# Differential effects of lysophospholipid headgroups, acyl chain length and saturation on vacuole acidification, Ca^2+^ transport, and fusion

**DOI:** 10.1101/2024.09.27.615487

**Authors:** Chi Zhang, Yilin Feng, Jorge D. Calderin, Adam Balutowski, Razeen Ahmed, Charlie Knapp, Ved Shah, Daniel Grudzien, Elizabeth Williamson, Jahnavi M. Karat, Rutilio A. Fratti

**Affiliations:** Deptartment of Biochemistry, University of Illinois Urbana-Champaign, Urbana, IL, 61801; Center for Biophysics and Quantitative Biology, University of Illinois Urbana-Champaign, Urbana, IL, 61801

**Author notes:** Correspondence: Rutilio A. Fratti.

**Keywords:** Acidification, Calcium, fusion, Lysophosphatidylcholine, Lysophosphatidic acid. SNARE, V-ATPase

## Abstract

SNARE-mediated membrane fusion is regulated by the lipid composition of the engaged bilayers. Lipids impact fusion through direct protein-lipid interactions or through modulating the physical properties of membranes to affect protein function. Lysophospholipids (LPLs) can affect membrane curvature, fluidity and energy of deformation. Their effects are due to their head group, and the length and saturation of their single acyl chains. Here we examined how the properties of LPLs affect yeast vacuole fusion and ion transport. We found that lysophosphatidylcholine (LPC) with acyl chains containing 14-18 carbons inhibited fusion with IC_50_ values of ≅ 40-120 µM. While acyl chain length moderately affected fusion, the head group played a major role. Unlike LPCs, Lysophosphatidic acid (LPA 18:1) failed to fully inhibit fusion, while lysophosphatidylethanolamine (LPE 18:1) had no effect. Separately we found that changes in acyl chain length and saturation differentially affected Ca^2+^ transport and vacuole acidification. Together these data show that the effects of LPLs on membrane fusion and ion transport were due to a combination of head group type and acyl chain length.

## 1 INTRODUCTION

Eukaryotes use SNARE-mediated membrane fusion to transport cargo between organelles as well as secretion at the plasma membrane (Bonifacino and Glick, 2004). Although the final stage of fusion is driven by SNAREs, the pathway is composed of multiple phases, each controlled by a distinct set of proteins and lipids. Membrane lipids can regulate protein function through two mechanisms. One mechanism is through specific interactions between lipids and proteins, as exemplified by the direct contacts of phosphoinositides (PIs) and lipid-binding protein domains (e.g., PH, PX, FYVE, etc.) (Lemmon, 2008; Balla, 2013; Overduin and Kervin, 2021). Another mode of regulation occurs through the effects of lipids on the physical properties of membranes including curvature, fluidity, thickness, and energy of deformation (Andersen and Koeppe, 2007; Harayama and Riezman, 2018; Levental and Lyman, 2023). These properties are affected by lipids such as cholesterol, sphingolipids, as well as glycerophospholipids with different acyl chain number, length and saturation, and head groups that vary in size and charge. Importantly, membrane curvature and fluidity can also be affected by lysophospholipids (LPL).

LPLs are generated through the action of various lipases and transferases. Lysophosphatidylcholine (LPC) can be generated by phospholipase-A1 (PLA1) or PLA2 on phosphatidylcholine (PC) to release the fatty acid at the sn-1 or sn-2 position, leaving an LPC with a single acyl chain and an inverted cone shape due to the large headgroup and narrow tail (Grzelczyk and Gendaszewska-Darmach, 2013). LPC can also be made by LCAT (lecithin-cholesterol acyltransferase) and transfer of acyl chain to cholesterol. LPC can be converted to PC by LPCAT (LPC acyl transferase) (Zhang et al., 2021). Lysophosphatidic acid (LPA) is made by the action of intracellular and extracellular autotaxin (LysoPLD) on LPC (Salgado-Polo et al., 2018; Xie et al., 2024). LPA can also be made through a two component pathway of soluble PLA2-IIA and LysoPLD (Aoki et al., 2002b; Tsuboi et al., 2015; Tserendavga et al., 2021). Lysophosphatidylserine (LPS) is made by a PS-specific PLA1 to make sn-2 LPS which converts to sn-1 (Aoki et al., 2002a, 2002a). Lysophosphatidylethanolamine (LPE) is made by PLA1/2 (Makiyama et al., 2025) and LPI is made by DDHD1, which was previously identified as a PA preferring PLA1 (Yamashita et al., 2013).

LPLs can function as signaling molecules that act through binding all four classes of human GPCRs to trigger signal transduction cascades (Radeff-Huang et al., 2004; Li et al., 2016; Shanbhag et al., 2020). LPLs can also insert into membranes to induce positive curvature, thinning, and affect fluidity by disordering lipid packing (Razinkov et al., 1998; Fuller and Rand, 2001; Mishima et al., 2004). The degree of curvature is dependent on the length and saturation the acyl chain and headgroup. The smallest head group is a single phosphate on LPA. Larger headgroups range from ethanolamine (LPE), serine (LPS), choline (LPC) to inositol (LPI). The acyl chains of LPLs can range from 10-24 carbons that are saturated or contain one or more double bonds (Shanbhag et al., 2020).

LPLs are mostly resistant to bilayer translocation, i.e. Flip-Flopping, which prevents equilibration between leaflets. LPLs with 12 carbon chains or shorter can flip-flop across membrane leaflets (Seeman, 1972; Lieber et al., 1984), whereas longer LPLs do not readily flip lead to a pronounced increase in positive curvature. In erythrocytes, LPC can be generated on either side of the plasma membrane causing distinct morphological features (Mohandas et al., 1978). LPC generation on the outer leaflet produces protrusions (echinocytes), whereas production on the inner leaflet is stomatocytogenic causing the formation of pits or invaginations on the cell surface. In both cases, the deformation is caused by positive curvature induced by local LPC production (Mohandas et al., 1978). The degree of positive curvature can influence membrane protein function as shown by the >100 fold difference in the dimerization of gramicidin channels when comparing LPC and LPI (Lundbaek and Andersen, 1994). Induced positive curvature can also promote membrane budding. Others have shown that ≥300 µM LPCs can produce bulges on 80 nm liposomes that bud off as 20-30 nm vesicles after long incubations (Arouri and Mouritsen, 2013; Hua et al., 2023). In yeast, LPI is enriched in COPII vesicles and promotes budding by increasing curvature and reducing membrane rigidity (Melero et al., 2018).

In addition to membrane deformation, LPLs can affect ion channels that play critical roles in homeostasis. For example, in erythrocytes LPA and LPCs regulate Ca^2+^/Mg^2+^ ATPase function in a manner dependent on acyl chain length and saturation, as well as headgroup type (Tokumura et al., 1985). In HEK-293 cells the transient receptor potential (TRP) Ca^2+^ channel TRPC5 is activated by LPC and LPI with acyl chains with ≥14 carbons, whereas LPL 12:0 has no effect (Flemming et al., 2006). Importantly, LPLs can affect Ca^2+^ transport in a protein-free manner through the formation of transient Ca^2+^ pores as seen in lymphoma cells (Wilson-Ashworth et al., 2004). LPLs can also trigger the opening of verapamil-sensitive L-type Ca^2+^ channels in monocytes (Lee et al., 2006). Other LPL effects include regulating HL60 differentiation to macrophages, guanylate/adenylate cyclase activity, and insulin receptor auto-phosphorylation (Shier et al., 1976; Asaoka et al., 1993; McCallum and Epand, 1995).

Altering membrane structure not only affects ion transport but can directly impact vesicle content mixing as seen with the fusion of liposomes with cells (Wu et al., 1996), microsome fusion (Chernomordik et al., 1993), or viral fusion (Chernomordik et al., 1997; Melikyan et al., 2000). Largely, this occurs through affecting the formation and stability of hemifusion diaphragms and the transition to full bilayer fusion (Melikyan et al., 1997). LPCs can also affect fusion at the priming stage when NSF activates SNAREs (Shin et al., 2012). The effect of LPLs on fusion can be compounded by palmitoyl or stearoyl LPC inhibition of PLC-dependent diacylglycerol (DAG) generation, which induces negative curvature needed for promoting the hemifusion to fusion transition (Okajima et al., 1998; Goñi and Alonso, 1999; Villar et al., 2000).

Yeast vacuole homotypic fusion serves as a model in vitro system to examine the role of specific proteins and lipids throughout the fusion cycle. Previous studies have mostly focused on glycerophospholipids, sphingolipids (SL), and ergosterol on the different stages of fusion (Kato and Wickner, 2001; Starr and Fratti, 2019; Zhang et al., 2024a). SNARE activation is regulated by the conversion of phosphatidic acid to DAG, as well as the presence of ergosterol and PI(4,5)P_2_ (Mayer et al., 2000; Kato and Wickner, 2001; Miner et al., 2016; Starr et al., 2019). Vacuole tethering is regulated by the Rab Ypt7 and HOPS complex, which are both controlled by phosphoinositides (PI) and SL (Stroupe et al., 2006; Cabrera et al., 2014; Lawrence et al., 2014; Zhang et al., 2024a). Tethering leads to docking and SNARE pairing at distinct membrane microdomains that form interdependently with PI3P, PI(4,5)P_2_, DAG, ergosterol and SL (Fratti et al., 2004; Zhang et al., 2024a). The formation of SNARE complexes triggers the release of Ca^2+^ from the vacuole lumen that is controlled in part by PI(3,5)P_2_ (Merz and Wickner, 2004; Miner et al., 2020). Prior to content mixing, vacuole outer leaflets undergo hemifusion which is affected by DAG, PI(3,5)P_2_, and LPC 12:0 (Reese and Mayer, 2005; Miner et al., 2019; Mondal et al., 2024). Between hemifusion and fusion others have shown that the integral membrane Vo complex of the V-ATPase can form dimers in trans between membranes in a LPC 12:0 sensitive manner (Peters et al., 2001).

## 2 Experimental procedures

### 2.1 Materials

Reagents were solubilized in PIPES-Sorbitol (PS) buffer (20 mM PIPES-KOH, pH 6.8, 200 mM sorbitol) with 125 mM KCl unless otherwise specified. Anti-Sec17 IgG (Mayer et al., 1996) and Pbi2 (Slusarewicz et al., 1997) were prepared as previously described. ATP was purchased from RPI (Mount Prospect, IL). Creatine phosphate was from Abcam (Waltham, MA). Acridine orange, Coenzyme A (CoA), Creatine kinase, Triton X-100 (TX-100) and FCCP (Carbonyl cyanide-4-(trifluoromethoxy) phenylhydrazone) were purchased from Sigma (St. Louis, MO) and dissolved in PS buffer or DMSO. Cal-520 dextran conjugate MW, 10,000 (Cal-520) was from AAT Bioquest (Sunnyvale, CA) and dissolved in DMSO. 1-myristoyl-2-hydroxy-sn-glycerol-3-phosphocholine (LPC 14:0), 1-palmitoyl-2-hydroxy-sn-glycero-3-phosphocholine (LPC 16:0), 1-stearoyl-2-hydroxy-sn-glycero-3-phosphocholine (LPC 18:0), 1-oleoyl-2-hydroxy-sn-glycero-3-phosphocholine (LPC 18:1), 1-oleoyl-2-hydroxy-sn-glycero-3-phosphatidic acid (sodium salt) (LPA 18:1), A23187, and CCCP (carbonyl cyanide m-chlorophenyl hydrazone) were purchased from Cayman Chemical (Ann Arbor, MI) and dissolved in PS buffer or ethanol. 1-oleoyl-2-hydroxy-sn-glycero-3-phosphatidylethanolamine (LPE 18:1) was from Avanti Polar Lipids (Birmingham, AL) and dissolved in ethanol.

### 2.2 Vacuole isolation and in vitro content mixing

Yeast vacuoles were isolated from BJ3505 (*pep4*Δ *PHO8)* and DKY6281 (*PEP4 pho8*Δ) as described with modifications (Jones et al., 1982; Klionsky and Emr, 1989; Haas et al., 1994). Content mixing reactions (30 μL) contained 3 μg each of vacuoles from BJ3505 and DKY6281 backgrounds in fusion reaction buffer composed of PS buffer, 125 mM KCl, 5 mM MgCl_2_, ATP regenerating system (ARS: 1 mM ATP, 0.1 mg/mL creatine kinase, 29 mM creatine phosphate), 10 μM CoA, and 280 nM recombinant Pbi2 (IB_2_, Protease B inhibitor) (Slusarewicz et al., 1997). The fusion of vacuoles mixes pro-Pho8 (inactive alkaline phosphatase) with the protease Pep4 to make mature Pho8. Reactions were incubated at 27°C for 90 min and Pho8 activity was assayed in 250 mM Tris-Cl, pH 8.5, 0.4% Triton X-100, 10 mM MgCl_2_, 1 mM *p*-nitrophenyl phosphate. Fusion was monitored as the absorbance at 400 nm from *p-*nitrophenolate production through phosphatase activity.

### 2.3 Fluorescence microscopy

Vacuole docking and vertex microdomain formation was performed as described (Sasser et al., 2012b). Isolated vacuoles harboring Vps33-GFP were treated with 100 µM LPC 14:0, 50 µM LPC 16:0, 18:0 and 18:1, 150 µM LPA or PS buffer alone under docking conditions (PS buffer, 100 mM KCl, 0.5 mM MgCl_2_, 0.33 mM ATP, 13 mM creatine phosphate, 33 µg/ml creatine kinase, 10 µM coenzyme A and 280 nM Pbi2) for 30 min at 27°C. After incubation, reactions were placed on ice and membranes were labeled with 4 µM FM4-64 (Wang et al., 2002). Reactions were next mixed with 20 µL 0.6% low melt agarose in PS buffer and mounted on microscope slides with cover slips. Images were taken with a Zeiss Axio Observer Z1 inverted microscope with an X-Cite 120XL light source, Plane Apochromat 63X oil objective (NA 1.4) and an AxioCam 705 mono R2 CCD camera. Images were analyzed and heat maps were generated with ImageJ software (NIH). Pseudo coloring and image merging was performed using Photoshop. s

### 2.4 Calcium transport

Ca^2+^ transport across the vacuole membrane was detected with Cal-520 dextran conjugate (MW 10,000) as described previously (Sasser et al., 2012a; Miner and Fratti, 2019; Miner et al., 2020). Ca^2+^ transport reactions (60 µl) contained 20 µg vacuoles isolated from BJ3505, reaction buffer, 10 µM CoA, 283 nM Pbi2, and 150 nM Cal-520. Reactions were transferred to wells of a black, half-volume 96-well flat-bottom plate with nonbinding surface. Reactions were incubated at 27°C for 5 min after which ARS was added to each well. For a negative control one well received additional PS buffer in lieu of ATP. Reactions were analyzed using a fluorescence plate reader (ex 485 nm and em 520 nm). SNARE-dependent Ca^2+^ efflux was blocked with antibody against the SNARE co-chaperone Sec17. This served as a negative control for SNARE function (Merz and Wickner, 2004). Fluorescence readings were taken every 30 s for 90 min and normalized to the initial fluorescence value set to 1 for each reaction.

### 2.5 Vacuole acidification

Vacuole acidification was monitored by acridine orange (AO) fluorescence as described previously (Müller et al., 2003; Zhang et al., 2022a, 2022b). Individual reactions (60 µl) contained 20 µg BJ3505 vacuoles, reaction buffer, ARS, 10 µM CoA, 283 nM Pbi2, and 15 µM AO. Reactions were set in a black half-volume 96-well flat-bottom plate with nonbinding surface. Test reagents or PS buffer was added, and reactions were incubated at 27°C for 60 s after which ARS was added to each well. One well received additional PS buffer in place of ATP to serve as a negative control. AO fluorescence (ex 485 nm; em 520 nm) was measured in a fluorescence plate reader and measurements were taken every 20 s. After AO fluorescence plateaued (400-600 s), each well received 30 µM FCCP to collapse the proton gradient and restore AO fluorescence. Each trace was normalized to their fluorescence intensities at the time of adding ATP and set to 1.

### 2.5 Statistical analysis

Results were expressed as the mean ± SE. Experimental replicates (n) are defined as the number of separate experiments. Statistical analysis was performed by One-Way ANOVA for multiple comparisons using Prism 10 (GraphPad, San Diego, CA). Tukey’s post hoc analysis was used for multiple comparisons and individual p-values. Statistical significance is represented as follows: \**p*<0.05, ** *p*<0.01, *** *p*<0.001, **** *p*<0.0001.

## 3 RESULTS

### 3.1 LPC and LPA inhibit vacuole fusion

Previous studies showed that vacuole fusion can be inhibited by LPCs with C10-14 saturated chains with IC_50_ values ranging from 30 µM for LPC 14:0 to ≅ 700 µM for LPC 10:0 (Reese and Mayer, 2005). Earlier work by Chernomordik et al. found that LPC 14:0 inhibited cell-cell fusion with and IC_50_ ≅ 10 µM and ≅ 1 µM for LPC 16:0 and LPC 18:0, respectively (Chernomordik et al., 1997). Together these studies showed the importance of tail length in altering fusion efficiency. Here we used LPCs containing C14-18 acyl chains. We also included the unsaturated LPC 18:1, LPA 18:1 and LPE 18:1 to compare monounsaturation as well as head groups. LPS was excluded due to the lack of solubility under fusion conditions. Vacuole fusion was determined by a content mixing assay (Jones et al., 1982; Klionsky and Emr, 1989).

In these experiments we added concentration curves of LPLs at the start of the reaction and incubated for 90 min at 27°C. We found that vacuole fusion was inhibited by LPC 14:0 with an IC_50_ of ≅ 120 µM **(Figure 1 A)**. When testing LPC 16:0 and LPC 18:0 we found that both inhibited fusion with IC_50_ values of ≅ 45 µM **(Figure 1 B-C)**. This was higher than the Chernomordik et al. study, however, a direct comparison is not possible due methodological differences and membrane types. Their assay monitored the hemagglutinin-dependent plasma membrane fusion of erythrocytes with HAb2 cells using R18 dequenching. This primarily measures outer leaflet fusion (i.e., hemifusion) and not necessarily full content mixing. Our work tracked the SNARE-dependent vacuole homotypic fusion using content mixing. Nevertheless, the level of fusion inhibition tracked with the increase in acyl chain length.

**Figure 1.**
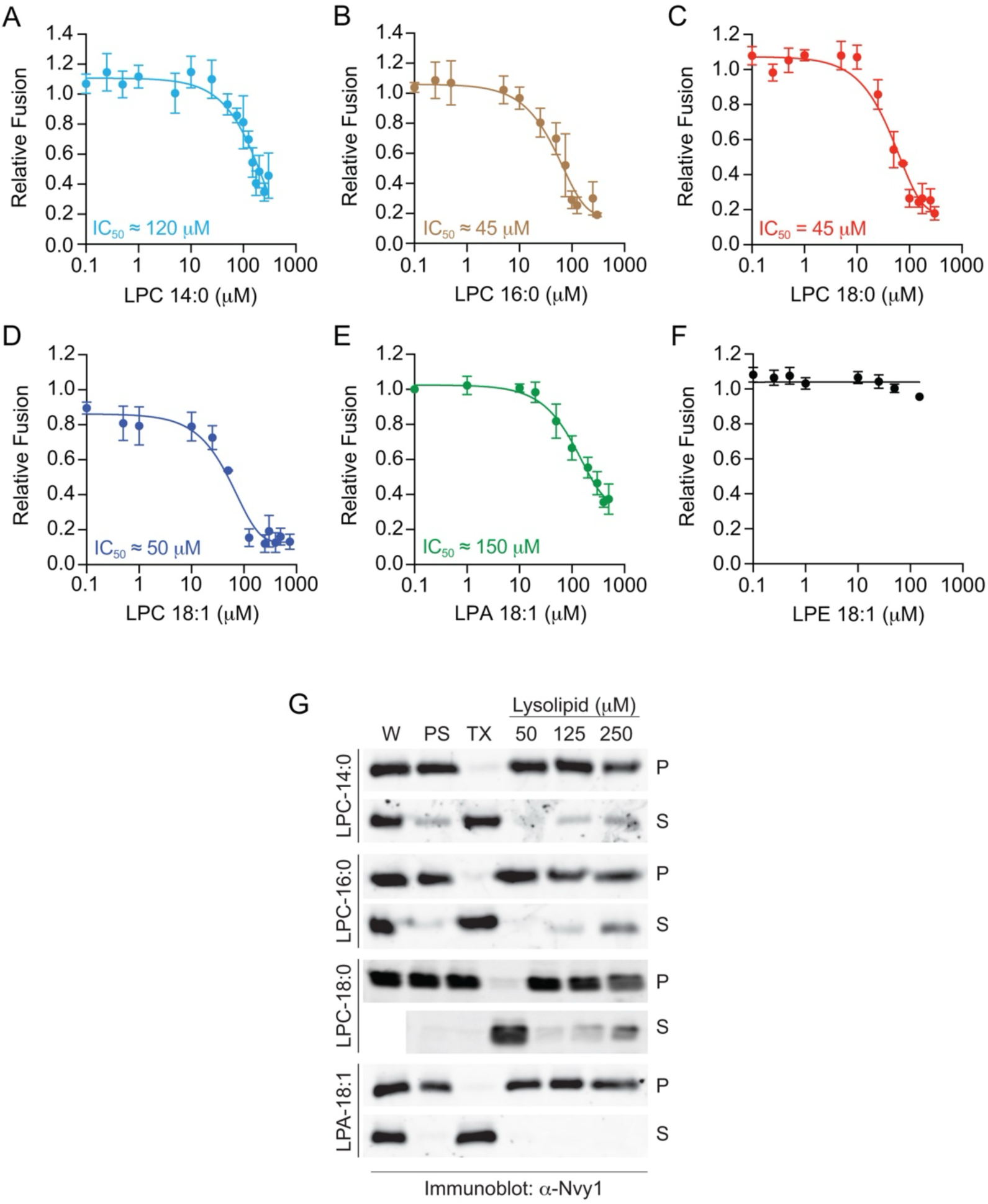
Inhibition of vacuole fusion by LPLs. Vacuole fusion reactions were treated with concentration curves of LPLs for 90 min at 27°C. Vacuole fusion reactions contained buffer or myristyl LPC 14:0 **(A)**, palmitoyl LPC 16:0 **(B),** stearoyl LPC 18:0 **(C)**, oleoyl LPC 18:1 **(D)**, oleoyl LPA 18:1 **(E),** or oleoyl LPE 18:1 **(F)**. Fusion values were normalized to the untreated control that is set to 1. Error bars represent mean ± SE. N≥3. **(G)** Purified vacuoles were incubated with PS buffer (PS), 1% Triton X-100 (TX), and LPL at the indicated concentrations. Reactions were incubated for 1 h at 30°C after which membranes were pelleted by centrifugation to separate vacuoles (P, pellet) from solubilized material (S, supernatant). Whole reactions (W), P and S were resolved by SDS-PAGE and immunoblotted for Nyv1.

We next asked if acyl chain saturation played a significant role in blocking fusion. To address this, we used oleoyl LPC 18:1 (*cis-*Δ^9^). Surprisingly, the unsaturation had no further effect as LPC 18:1 inhibited fusion with an IC_50_ ≅ 50 µM **(Figure 1 D)**. Finally, we examined the role of headgroup type by comparing LPC 18:1 with LPA 18:1 and LPE 18:1. This showed that LPA affected vacuole fusion but required a much higher concentration with an IC_50_ of ≅ 150 µM, which is an underestimate since inhibition was only reduced by 60% **(Figure 1 E)**. Finally, we found that LPE 18:1 had no effect on fusion **(Figure 1 F)**. This suggested that the head group of LPLs was more impactful on fusion versus acyl chain length. The differences could be due to variances in induced positive curvature. Both LPC and LPA are known to induce positive curvature, whereas LPE has no effect on curvature (Fuller and Rand, 2001; Kooijman et al., 2005).

### 3.2 LPC and LPA do not lyse vacuoles at levels that block fusion

Because LPLs can have detergent effects we tested if the inhibition of fusion vacuoles was due to lysis. We used LPLs at their approximate IC_50_ values as well as concentrations that exceeded these values. To probe for lysis, we pelleted membranes after the fusion reaction was complete and collected the supernatant. Both membrane and supernatant fractions were probed by Western blotting for the membrane anchored SNARE Nyv1. As a control for lysis, we used 1% TX-100. In **Figure 1 G** we show that under control (PS) conditions Nyv1 was only in the membrane/pellet (P) fraction apart from some background signal due to incomplete membrane re-isolation as well as non-specific lysis during fusion (Starai et al., 2007). In contrast, Nyv1 was shifted to the supernatant when vacuoles were treated with TX-100. No LPC variant lysed vacuoles at 50 and 125 µM. Nyv1 was only solubilized when LPC 16:0 and LPC 18:0 concentrations were at 250 µM which was five-fold higher than their IC_50_ values. LPA had no effect on Nyv1 solubilization at any concentration tested. Together, these data indicate that fusion inhibition was not due to lytic effects.

LPLs can act as detergents; however, this activity varies widely depending on the type of membrane (liposomes, purified organelles, whole cells, etc.), lipid composition, phase separation, and duration of exposure. For instance, LPC 18:0 can lyse microsomes but requires ≥400 µM, whereas erythrocytes are lysed in seconds by <20 µM LPCs (Weltzien, 1979; Tanaka et al., 1983; Chernomordik et al., 1993). Protein-free liposomes can be partially lysed by LPC with different acyl chains and dependent on osmolarity (Ralston et al., 1980; Senisterra et al., 1988). The ability of LPLs to permeabilize membranes can be counteracted by cholesterol, which in turn explains some of the differences between the susceptibility of artificial liposomes versus biological membranes (Arouri and Mouritsen, 2013). Another reason for such differences could be linked to the initial curvature of the experimental membrane so that smaller vesicles with high curvature are less disturbed by LPLs (Jespersen et al., 2012). For instance, fragile erythrocytes are 6-8 µm wide, while more resistant microsomes are 20-200 nm across and liposomes range from 50 nm to 100 µm. When considering true detergents, solubilization requires crossing bilayers to act on both leaflets, and full micellar solubilization can take hours (Kragh-Hansen et al., 1998; Hua et al., 2023).

### 3.3 LPLs do not affect vertex microdomain formation

A study by Reese and Mayer showed that LPCs inhibited fusion after docking with moderate effects on trans-SNARE complex formation (Reese and Mayer, 2005). Thus, prior stages such as priming and docking were likely unaffected. That said, LPLs have been shown to alter microdomain size suggesting that vacuolar microdomains could be altered (Krasnobaev et al., 2022). During vacuole docking the lipids and proteins that drive fusion accumulate into vertex membrane microdomains and alterations in their assembly can inhibit downstream protein function (Wang et al., 2003; Fratti et al., 2004; Sasser et al., 2012b; Zhang et al., 2024b). Here we tested if LPLs at their IC_50_ concentrations altered vertex microdomain assembly. As a reporter we used vacuoles that contained Vps33-GFP, a subunit of the HOPS tethering complex (Wang et al., 2003). The vital dye FM4-64 was used to label the entirety of the docked membranes. Images were overlaid/merged to show where GFP and FM4-64 colocalized and heat maps were generated to show the intensities of GFP localization in relation to the other parts of the docked vacuoles. We found that Vps33-GFP enrichment at vertices was not affected by any of LPC or by LPA **(Figure 2)**. This was consistent with the previous study showing that the effect of LPLs on fusion was downstream of membrane docking.

**Figure 2.**
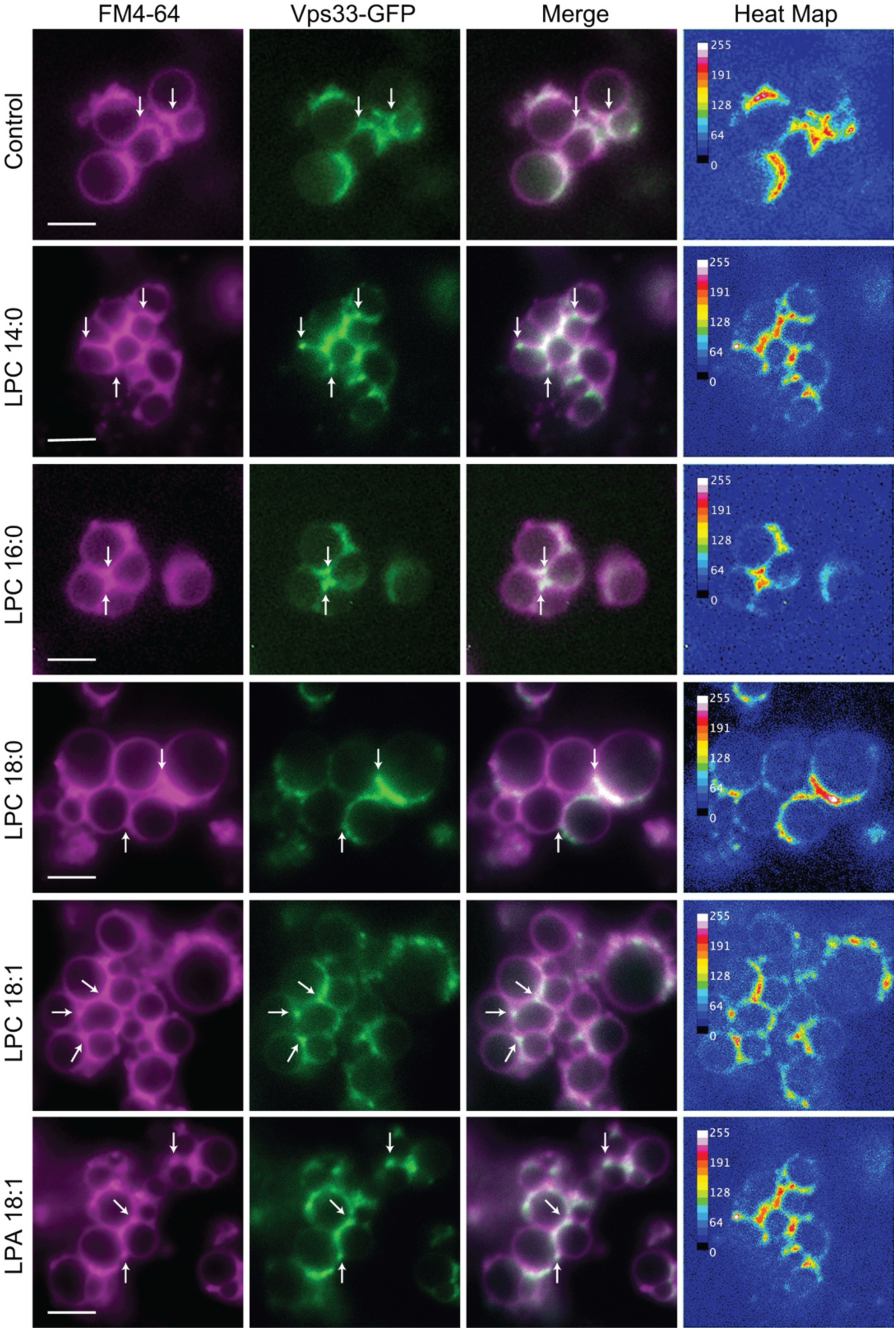
Purified vacuoles containing Vps33-GFP were treated with 50 µM LPC 16:0, LPC 18:0 and LPC 18:1, 150 µM LPC 14:0, 150 µM LPA 18:1 or buffer alone (control) and incubated for 30 min at 27°C. Limiting membranes were labeled with FM4-64. Vps33-GFP labeled vertex microdomains. Shown are individual channels, merged images and heat maps to show GFP intensities. Arrows show vertex microdomain examples. Scale bar, 4 µm.

### 3.4 Ca^2+^ transport across the vacuole membrane is sensitive to LPLs

Vacuoles are the main Ca^2+^ storage organelle in yeast, containing mM levels of the cation that is mostly complexed to polyphosphate, leaving µM free Ca^2+^ available for transport (Dunn et al., 1994). Vacuolar Ca^2+^ uptake utilizes the high affinity Ca-ATPase pump Pmc1 and the low affinity Ca^2+^/H^+^ (K^+^/H^+^) exchanger Vcx1 (Cunningham and Fink, 1994, 1996). Vacuoles also contain the PI(3,5)P_2_ sensitive TRP homolog Yvc1 that releases Ca^2+^ upon hyperosmotic stress-induced vacuole fission (Bonangelino et al., 2002; Denis and Cyert, 2002; Dong et al., 2010; Su et al., 2011).

During vacuole fusion SNAREs form complexes between membranes that leads to the release of luminal Ca^2+^ (Merz and Wickner, 2004). This mechanism is independent of Yvc1, yet Ca^2+^ release remains sensitive to levels of PI(3,5)P_2_ (Miner et al., 2020). Since Ca^2+^ release during fusion was likely mechanosensitive we next tested the effects of LPLs with a real-time fluorescence assay to measure the kinetics of Ca^2+^ uptake and efflux (Sasser et al., 2012a; Miner and Fratti, 2019). Extraluminal Ca^2+^ was detected with membrane impermeable Cal-520-dextran, which only fluoresces when bound to Ca^2+^. Cal-520 fluorescence was high at the start of the reaction and rapidly decreased after adding ATP to activate Ca^2+^ uptake. Upon trans-SNARE complex formation, Ca^2+^ was released leading to an increase in fluorescence. As a control for efflux we included anti-Sec17 IgG to prevent SNARE activation (Mayer et al., 1996). This allowed Ca^2+^ uptake without a subsequent release. As a negative control for uptake, we included a reaction that lacked ATP resulting in steady Cal-520 fluorescence.

To test the effects of LPLs on Ca^2+^ transport we added them at the start of the reactions, before adding ATP to test their effects on uptake. Starting with LPC 14:0, we observed reduced Ca^2+^ uptake as shown by the partial loss of fluorescence that stopped at 10 min and leveled off, thus masking any effect on efflux at 30 min **(Figure 3 A-C)**. Based on the fusion data we imagined that LPC 16:0 would have a stronger effect on Ca^2+^ uptake versus LPC 14:0. On the contrary, LPC 16:0 had no effect on Ca^2+^ uptake and resembled the untreated control **(Figure 3 D-F)**. While uptake was not affected, LPC 16:0 completed blocked Ca^2+^ efflux at 125 µM and 250 µM and partially blocked at 50 µM. When testing LPC-18:0 we found that it inhibited uptake to a greater extent compared to LPC 14:0 **(Figure 3 G-I)**. The uptake of Ca^2+^ was inhibited by 250 µM LPC 18:0, while 125 µM kept uptake from reaching the anti-Sec17 plateau indicating that uptake was partially blocked. The effect of 250 µM LPC 18:0 might be due to lysis whereas the effect of 125 µM LPC 18:0 likely masked any change in efflux. LPC 18:0 at 50 µM had no effect on uptake but did cause a delay in efflux.

**Figure 3.**
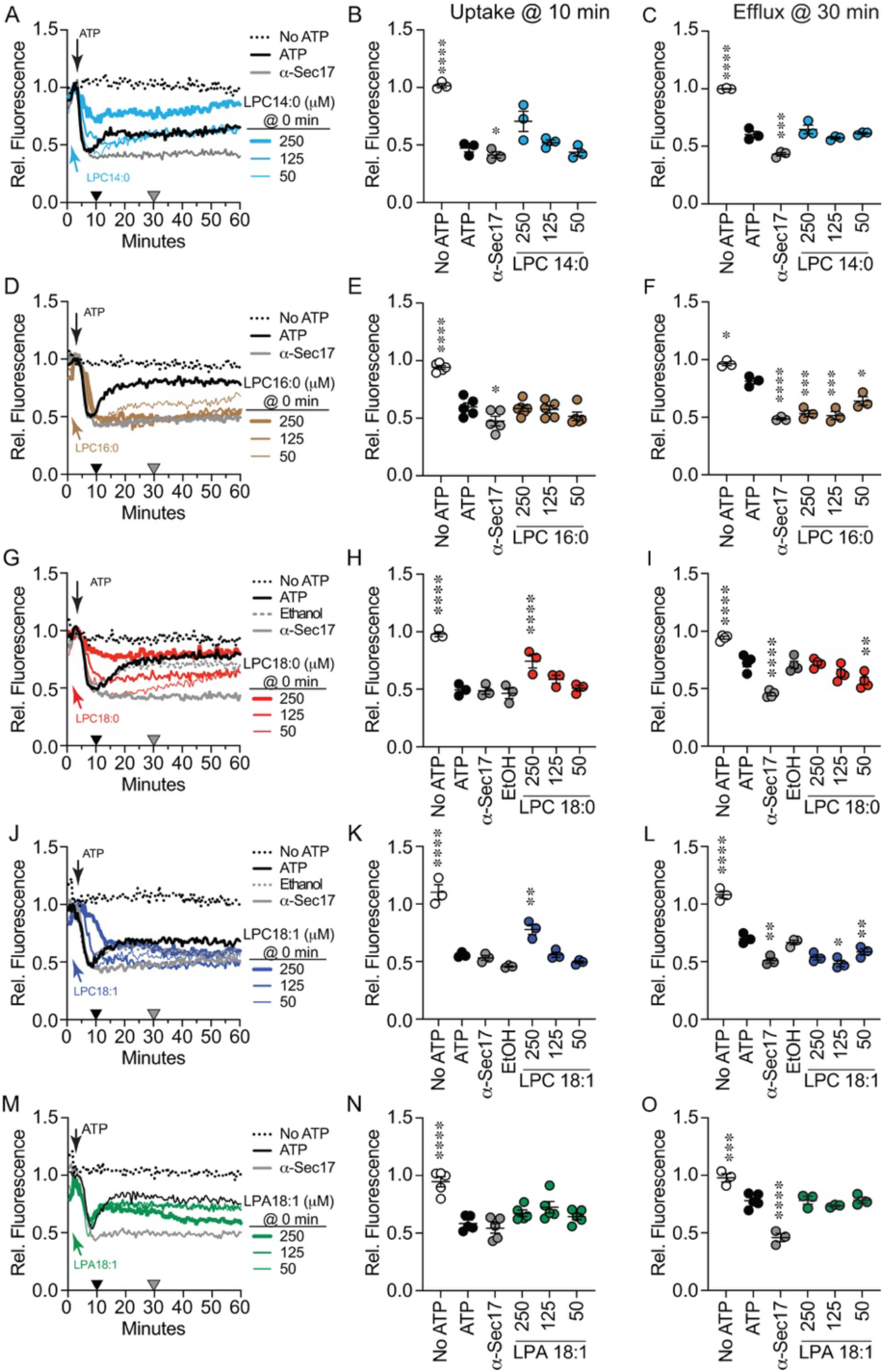
Effect of LPLs on Ca^2+^ uptake. Vacuoles were isolated from BJ3505 and 2X fusion reactions (60 µL) were treated with 140 µg/mL α-Sec17 IgG, LPC variants or reaction buffer at T=0 min in the absence of ARS and in the presence of Cal-520. Reactions contained LPC 14:0 **(A)**, LPC 16:0 **(D)**, LPC 18:0 **(G)**, LPC 18:1 **(J)**, or LPA 18:1 **(M)**. After 5 min of incubation ARS or buffer was added to each reaction. Cal-520 fluorescence was monitored every 30 s for 60 min. Fluorescence values were normalized to the untreated control at T=5 when ATP was added and set at 1. **(B, E, H, K, N)** Ca^2+^ uptake efficiency was quantified after 10 min of incubation (Black triangles). **(C, F, I, L, O)** SNARE-dependent Ca^2+^ efflux efficiency was quantified after 30 min incubation (Gray triangles). Data points represent mean ± SE (n≥3). Significance was measured using one way ANOVA for multiple comparisons. Tukey’s post hoc test was used for individual p values. \*\**p*<0.01, \*\*\**p*<0.001, \*\*\*\**p*<0.0001.

Next, we asked whether acyl chain saturation and head group affected Ca^2+^ transport. First, we tested LPC 18:1 and found that it delayed and reduced uptake at 250 µM, but to a lesser extent versus LPC 18:0 **(Figure 3 J-K)**. At lower concentrations LPC 18:1 did not significantly alter uptake, however Ca^2+^ efflux at 30 min was inhibited **(Figure 3 J & L)**. This suggested that a monounsaturation reduced the block of uptake while minimally changing the inhibition of efflux. Finally, we asked if headgroup type would affect Ca^2+^ transport. To answer this, we used LPA 18:1 and found that it had no effect on Ca^2+^ uptake or efflux when added at the beginning of the experiment **(Figure 3 M-N)**. This indicated that the type of the head group played a critical role in altering Ca^2+^ flux.

The effects of LPLs on Ca^2+^ uptake were not due to membrane permeabilization. In separate experiments we tested the effects of TX-100 and the Ca^2+^ ionophore A23187. Adding TX-100 or A23187 at the start of the reaction completely blocked Ca^2+^ uptake as measured by the lack of fluorescence quenching of Cal-520 **(Supplemental Figure 1)**.

In previous studies we found that effects on Ca^2+^ uptake and efflux can be separated by varying when an inhibitor is added (Miner et al., 2020). In Figure 2 we added LPLs at the start of the assay and monitored both uptake at 10 min and efflux at 30 min. Some of the LPLs blocked uptake, masking any effect on efflux later in the pathway. We next tested if LPLs could alter efflux when added at 10 min after uptake was completed. Adding LPC 14:0 at 10 min led to a brief Ca^2+^ release that was quickly reabsorbed **(Figure 4 A-B)**. We also observed delayed Ca^2+^ efflux for ∼10 min after which the reactions rapidly recovered to match the untreated control at 30 min **(Figure 4 A-B)**. LPC 16:0 added at 10 min also caused a shortened efflux followed by a rapid reuptake **(Figure 3C-D)**. However, unlike LPC 14:0, LPC 16:0 blocked Ca^2+^ efflux at the concentrations tested. Interestingly, LPC 18:0 had no striking effect on efflux even though the effect of 50 µM was statistically significant **(Figure 4 E-F)**. This showed that LPC 18:0 affected uptake but not efflux. Together these data indicated that the most effective acyl chain length for altering Ca^2+^ efflux was 16 carbons. Since IC_50_ values for inhibiting fusion were identical for LPC 16:0 and - 18:0, the difference in altering Ca^2+^ transport suggested that there was an additional effect triggered by LPC 16:0 to alter the pathway.

**Figure 4.**
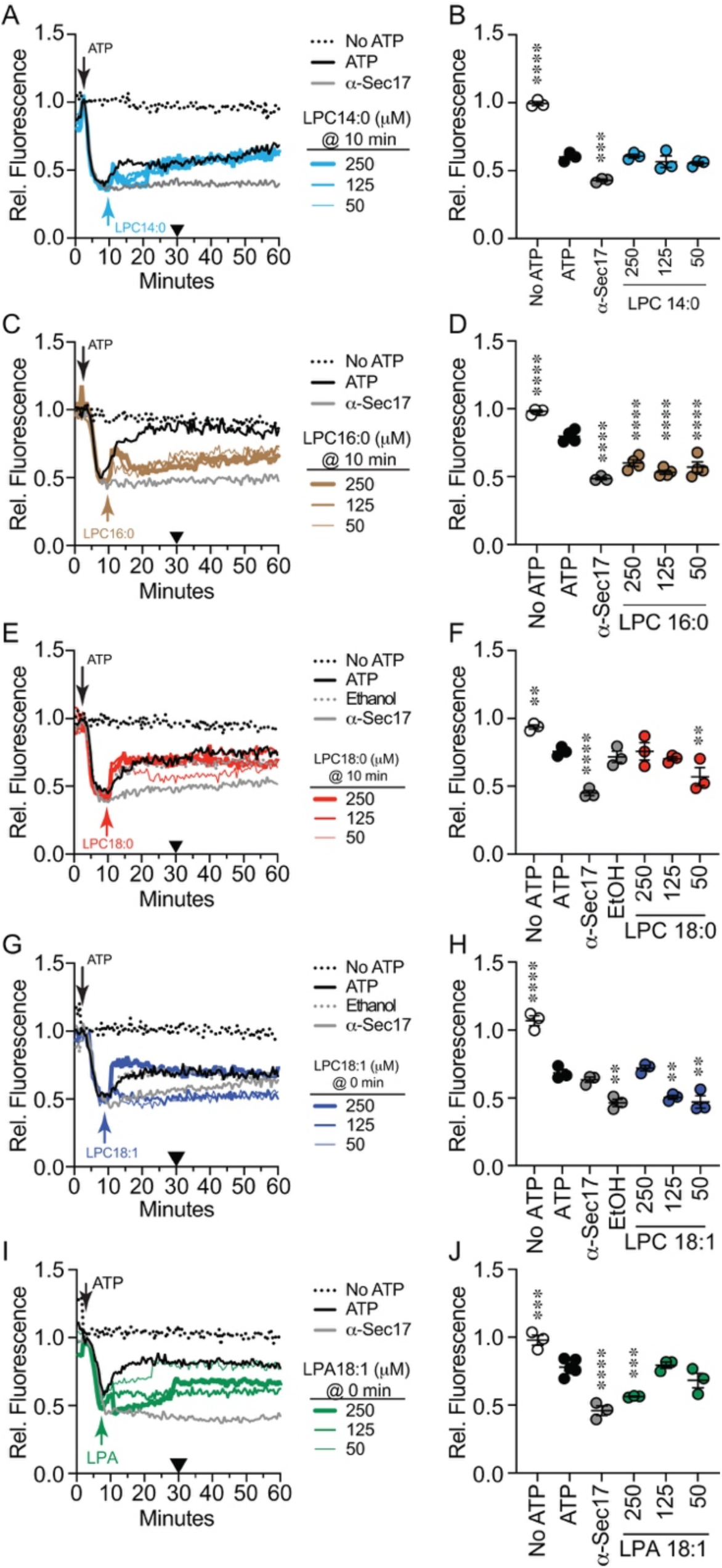
Effect of LPLs on Ca^2+^ efflux. Vacuoles from BJ3505 were incubated as 2X fusion reactions (60 µL) and treated with 140 µg/mL α-Sec17 IgG at T=0 in the presence of Cal-520. After 5 min of incubation ATP or buffer was added to each reaction. LPC variants were added after 10 min. Reactions contained LPC 14:0 **(A)**, LPC 16:0 **(C)**, LPC 18:0 **(E)**, LPC 18:1 **(G)**, or LPA 18:1 **(I)**. Cal-520 fluorescence was monitored every 30 s for 60 min. Fluorescence values were normalized to the untreated control at T=5 min when ARS was added and set to 1. **(B, D, F, H, J)** SNARE-dependent Ca^2+^ efflux efficiency was quantified after 30 min incubation (Black triangles). Data points represent mean ± SE (n≥3). Significance was measured using one way ANOVA for multiple comparisons. Tukey’s post hoc test was used for individual p values. \**p*<0.05, \*\**p*<0.01, \*\*\**p*<0.001, \*\*\*\**p*<0.0001.

Next, we tested if acyl chain saturation or head group size affected Ca^2+^ efflux. When added after Ca^2+^ uptake was complete we found that 250 µM LPC 18:1 triggered a rapid rise in efflux **(Figure 4 G-H)**. This was likely due to partial lysis. After the rapid rise, the rate leveled off to match the untreated control. Importantly, lower levels of LPC 18:1 potently inhibited Ca^2+^ efflux.

Lastly, we tested the late addition of LPA 18:1 on Ca^2+^ transport. When added at the start LPA had no effect on uptake or efflux; however, when added after uptake was complete, we observed a dose dependent inhibition of Ca^2+^ efflux **(Figure 4 I-J)**. This difference could be due to the extraluminal Ca^2+^ concentration at the time of addition. For instance LPA forms type-1 lipid micelles in the absence of divalent cations whereas the presence of divalent cations drives LPA to form complexes with the cation in a bilayer (Kooijman et al., 2003).

Again, the inhibitory effects of LPLs on Ca^2+^ transport were not due to membrane permeabilization. Ca^2+^ efflux was tested in the presence of TX-100 and A23187 added after Ca^2+^ uptake was completed. Both TX-100 or A23187 caused a near-instant release of Ca^2+^ characterized by a near vertical spike in Cal-520 fluorescence **(Supplemental Figure 1 A-B)**. To better observe the signal spike, we added TX-100 and A23187 in the presence of anti-Sec17 to block natural SNARE dependent Ca^2+^ efflux. This had no effect on Ca^2+^ release triggered by TX-100 and A23187 **(Supplemental Figure 1 C-D)**.

### 3.5 Vacuole acidification is inhibited by LPC

Yeast vacuoles are acidic lysosomal compartments that aid in the degradation of macromolecules into their constituent building blocks. This requires an acidic environment that activates digestive enzymes at the site of activity and not during their trafficking through other organelles. Vacuole acidification is primarily carried out by the vacuolar ATPase (V-ATPase) that is composed of two complexes. The V_O_ complex is embedded in the membrane whereas the V_1_ complex is cytoplasmic and can reversibly associate with the V_O_ to form the active holoenzyme (Vasanthakumar and Rubinstein, 2020). The stability of V_1_-V_O_ is promoted in part through the interactions of the V_O_ subunit Vph1 and PI(3,5)P_2_ (Li et al., 2014; Banerjee et al., 2019). Additionally, vacuole acidification is attenuated by the lack of SL which increases fluidity and inhibits vacuole fusion (Zhang et al., 2024a). The activity of V-ATPases is also sensitive to the fluidizing anesthetic dibucaine. Aside from fluidity, we have found that acidification is sensitive to high Ca^2+^ (Zhang et al., 2022b). Based on this and our data showing that LPLs can affect Ca^2+^ transport, we next asked if they could affect vacuole acidification.

To measure vacuole acidification, we used AO which shifts fluorescence emission in acidic environments (Moriyama et al., 1982). AO fluorescence shifts from 520 nm to 680 nm when transported in to the vacuole lumen (Pierzyńska-Mach et al., 2014; Thomé et al., 2016; Zhang et al., 2022a). To test the effects of LPLs on acidification reactions were with run with buffer alone or different concentrations of LPL for 30 sec after which ATP was added to start V-ATPase activity. A separate reaction that lacked ATP served as a negative control. Reactions were run for 600 s at which point 30 µM of the protonophore FCCP was added to each well to equilibrate H^+^ concentrations across the membrane and restore AO fluorescence at 520 nm. This showed that the decrease in fluorescence was indeed due to a proton gradient. In **Figure 5A** the black trace shows the positive control with ATP and no LPC. The decrease in fluorescence plateaued at ∼400 s after the addition of ATP. In the absence of ATP, AO fluorescence was maintained throughout the incubation (dotted trace). The addition of LPC 14:0 decreased the rate of acidification in a dose dependent manner **(Figure 5 A-B)**. That said, each concentration of LPC 14:0 caught up with the untreated control by 600 s. When we measured fluorescence at 400 s for multiple experiments, we found that LPC 14:0 significantly attenuated H^+^ transport (**Figure 5 B)**. This proportionally matched its effect on Ca^2+^ uptake when added at T=0.

**Figure 5.**
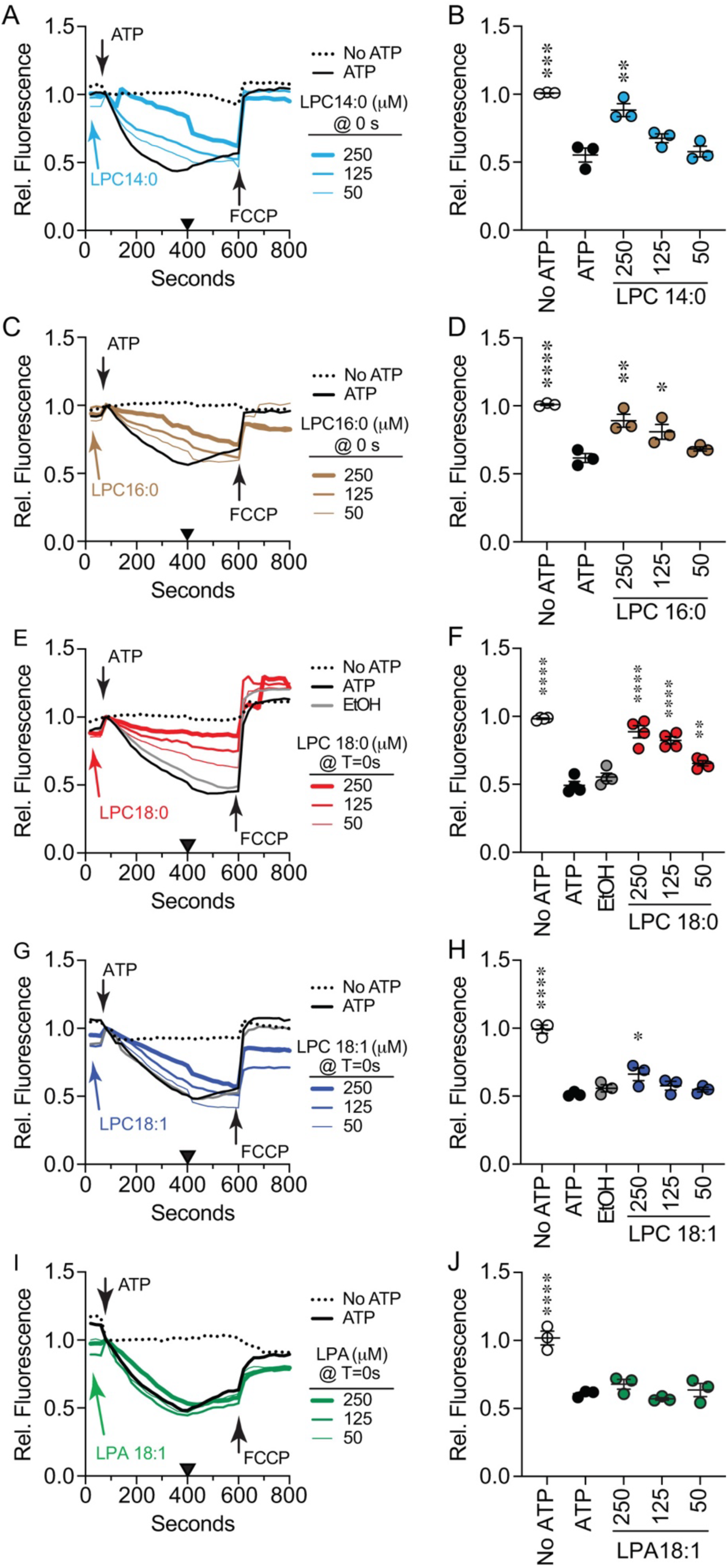
Effect of LPLs on initial vacuole acidification. BJ3505 vacuoles fusion reactions (2X) were incubated with LPC variants or reaction buffer at T=0 min in the absence of ARS and in the presence of AO. After 60 s of incubation received ARS or additional buffer and further incubated for a total of 800 s. After 600 s each reaction received FCCP to equilibrate the proton gradient. AO fluorescence was normalized to the fluoresced at the time of adding ARS and set to 1. Separate experiments were treated with LPC 14:0 **(A)**, LPC 16:0 **(C)**, LPC 18:0 **(E)**, LPC 18:1 **(G)**, or LPA-18:1 **(I). (B, D, F, H, J)** Vacuole acidification efficiency was quantified after 400 s (Black triangles). Data ponts represent mean ± SE (n≥3). Significance was measured using one way ANOVA for multiple comparisons. Tukey’s post hoc test was used for individual p values. \**p*<0.05, \*\**p*<0.01, \*\*\**p*<0.001, \*\*\*\**p*<0.0001.

We next tested the effects of longer acyl chains on V-ATPase activity. Like LPC 14:0, the addition of LPC 16:0 transiently slowed V-ATPase activity in a dose dependent manner that caught up with the control by 600 s **(Figure 5 C-D)**. Interestingly, treating reactions with LPC 18:0 appeared to stably inhibit V-ATPase activity as the treated samples never caught up with the control **(Figure 5 E-F)**. Together these data indicated that the V-ATPase was sensitive to acyl chain length. To length alone was responsible for the effect, we used oleoyl LPC 18:1. We found that LPC 18:1 only slowed V-ATPase activity, and the treated samples caught up with the control as seen with the with the shorter acyl chain containing LPCs **(Figure 5 G-H)**. This suggested that the narrowing of stearoyl vs oleoyl hydrocarbons influenced the inhibition of V-ATPase activity. Lastly, we tested LPA 18:1 and observed that vacuole acidification was unaffected **(Figure 5 I-J)**. This showed that an oleoyl acyl chain alone was not sufficient in altering vacuole acidification. Instead, it appeared that inhibiting V-ATPase function with LPLs was linked to the length of a saturated acyl chain and the head group.

Finally, we examined the effect of adding LPLs after vacuole acidification had plateaued. We found that LPC 14:0 had no significant effect on maintaining acidification levels **(Figure 6 A-B)**. In contrast, LPC 16:0 **(Figure 6 C-D)**, LPC 18:0 **(Figure 6 E-F)** and LPC 18:1 **(Figure 6 G-H)** all reversed V-ATPase activity as shown by the rise in AO fluorescence, albeit to different degrees. This suggested that once acidified, maintaining the H^+^ gradient was sensitive to acyl chains of 16 or more carbons in length and independent of hydrocarbon mono-unsaturation. Importantly, LPA 18:1 had no effect when added late matching its early addition **(Figure 6 I-J)**. This further indicated that head group type was critical in the inhibition of vacuole acidification.

**Figure 6.**
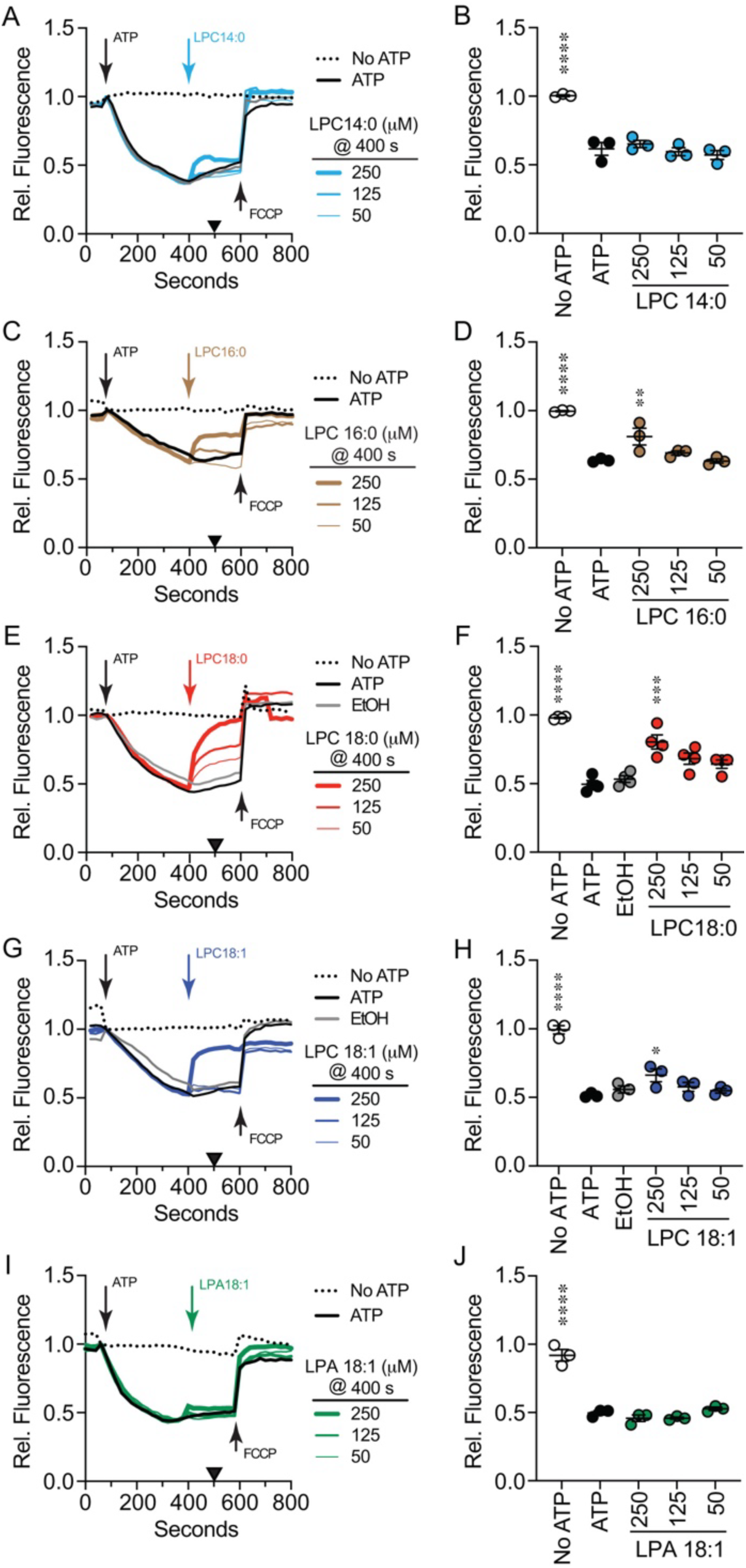
Effect of LPLs on maintaining vacuole acidification. Vacuoles from BJ3505 were incubated as 2X fusion reactions (60 µL) and incubated with reaction buffer and AO at T=0 min. After 60 s of reactions received ARS or additional reaction buffer. LPC variants were added at 400 s. After 600 s each reaction received FCCP to equilibrate the proton gradient and further incubated for a total of 800 s. AO fluorescence was normalized to the fluoresced at the time of adding ARS and set to 1. Separate experiments were treated with LPC 14:0 **(A)**, LPC 16:0 **(C)**, LPC 18:0 **(E)**, LPC 18:1 **(G)**, or LPA 18:1 **(I). (B, D, F, H, J)** Vacuole acidification efficiency was quantified after 400 s (Black triangles). Data points represent mean ± SE (n≥3). Significance was measured using one way ANOVA for multiple comparisons. Tukey’s post hoc test was used for individual p values. \*\*\**p*<0.001, \*\*\*\**p*<0.0001.

Similar to the effects of LPLs on Ca^2+^ transport, their effects on vacuole acidification were not linked to lysis. In separate experiments we used TX-100 and different protonophore CCCP to test acidification. As expected, the addition of either TX-100 or CCCP at the start of the reaction fully blocked acidification **(Supplemental Figure 2)**. We also tested the effects of TX-100 and CCCP after vacuole acidification was complete. For both reagents we observed a near vertical spike in AO fluorescence, thus demonstrating that the effects of LPLs on acidification were not due to lysis.

## 4 DISCUSSION

In this study we examined the role of LPLs on vacuole fusion, Ca^2+^ transport and V-ATPase activity. Specifically, we tested the roles of acyl chain length and monounsaturation as well as headgroup type. In general, we found that the LPL traits tested had different effects on fusion and ion transport. Apart from LPE, each of the LPLs inhibited vacuole fusion with varying efficacy. For fusion the critical factor was headgroup type with LPA being less effective relative to the LPC variants tested. The dependence on headgroup type suggested that differences in blocking fusion could be due to the degree of positive curvature. While acyl chain length and saturation can further affect curvature, it did not play a large role in blocking fusion. To illustrate the impact of head group type, oleoyl LPC has a spontaneous positive curvature radius of +38 Å versus oleoyl LPA with a radius of +20 Å in DOPE liposomes (Fuller and Rand, 2001; Kooijman et al., 2005). Palmitoyl LPC has a curvature radius of +68 Å, illustrating how acyl chain saturation can further influence membrane remodeling. On the other hand, LPE does not alter curvature in model membranes. In addition to changes in curvature, it is possible that the negative charge of LPA influences how well fusion is blocked. For instance, the negative charge of LPA can interact with Ca^2+^ to reduce its effect on curvature by inducing bilayer formation (Kooijman et al., 2003).

While the acyl chain length of 14-18 carbons made little difference in how well LPCs inhibited fusion a study by Reese and Mayer looked at shorter acyl chain variants and found that inhibition was relative to hydrocarbon length (Reese and Mayer, 2005). Specifically, they found that LPC-14, −12, −10 and −8 inhibited vacuole fusion with IC_50_ values of 30 µM, 100 µM, 400 µM, and 3 mM, respectively (Reese and Mayer, 2005). Together with our data, we can conclude that acyl chain length is important until efficacy plateaus at 14C. Their study went on to show that the block in fusion was between SNARE pairing/hemifusion and content mixing. A critical step between these phases is the release of luminal Ca^2+^ upon trans-SNARE complex formation (Merz and Wickner, 2004). The effects of LPLs on Ca^2+^ transport have been shown with multiple transporters in different cell types, however, these studies mostly focused on the plasma membrane and cytosolic Ca^2+^.

Among the types of Ca^2+^ transporters affected by LPLs are L-type voltage dependent channels. In myocytes production of LPC stimulates L-type channels leading to Ca^2+^ overloading that was inhibited by the L-type antagonist nimodipine (Golfman et al., 1999). The same is seen in Drosophila S2 cells (Wang et al., 2015). This effect was only seen with LPC 18:0 and LPC 16:0, whereas LPC 18:1, −18:2, −18:3 had no effect. Furthermore, LPE, LPA, LPI and LPS did not trigger Ca^2+^ accumulation. Transport through L/P-type channels including the yeast Ca^2+^ P-type ATPase are sensitive to channel blockers such as verapamil (Zhang et al., 2022b).

LPLs can also affect Ca^2+^ transport by mechanosensitive channels including members of the TRP family. For example, TRPC5 is activated by LPC and LPI with acyl chains containing 14 or more carbons (Flemming et al., 2006). That said the same study showed that LPLs had no effect on TRPC2. LPC and LPI also induce Ca^2+^ influx via TRPv2 when the acyl chains were at least 16 carbons in length. This also led to increased cell migration in prostate cancer cells, suggesting that actin dynamics were affected (Monet et al., 2009). In this case, headgroup type was also considered and revealed that LPA and LPE had no effect. The effect of headgroup type is not consistent as shown in lens cells where LPA triggers Ca^2+^ flux through a mechanosensitive channel, while LPC had no effect (Ohata et al., 1997). In these cases, LPL triggered transport through mechanosensitive ion channels are not affected by inhibitors such as verapamil and nicardipine, thapsigargin or blocking phospholipase-C with U73122.

So, how do LPLs affect yeast vacuole Ca^2+^ transport? Vacuoles move Ca^2+^ from the cytoplasm to the lumen through the P-type ATPase Pmc1 and the Ca^2+^/H^+^ antiporter Vcx1. Previously, we showed that Ca^2+^ uptake is blocked by verapamil when added at T=0, which looks like the effects of LPC 14:0 and LPC 18:0 and to a lesser extent LPC 18:1. On the other hand, LPC 16:0 and LPA 18:1 have no effect on uptake whereas LPC 16:0 completely blocks Ca^2+^ efflux. This also eliminates the possibility of causing lysis or transient pores to release Ca^2+^ as seen in lymphoma cells (Wilson-Ashworth et al., 2004). When added late in the reaction we previously showed that verapamil caused a rapid release of Ca^2+^, suggesting that Pmc1 activity normally continues throughout the reaction in part to balance the release of Ca^2+^. Otherwise, only LPC 16:0 and LPA 18:1 were able to block Ca^2+^ efflux. LPC 18:1 blocked Ca^2+^ release at 50 and 125 µM whereas 250 µM triggered a rapid release. The later could be attributed to lysis whereas the former indicates that efflux was truly inhibited.

Regarding acidification, we believe this is likely occurring through changes in curvature and/or fluidity. V-ATPase activity has been linked to its interactions with cholesterol-rich microdomains in tumor cells (Costa et al., 2018). Depletion of cholesterol with methyl-β-cyclodextrin, therfore increasing fluidity reduced pumping activity as well as rates of cell migration. In our hands, increasing fluidity with dibucaine or SL attenuates V-ATPase function (Zhang et al., 2024a).

Could LPLs interdependently affect Ca^2+^ and H^+^ transport? We previously showed that vacuole acidification was interdependent with Ca^2+^ gradients (Zhang et al., 2022b). The addition of Ca^2+^ to AO assays inhibits V-ATPase function that is dependent on the Ca^2+^/H^+^ exchanger Vcx1 (Zhang et al., 2022b). Conversely, Ca^2+^ transport required vacuole acidification. Both chloroquine and the V-ATPase inhibitor Bafilomycin block Ca^2+^ uptake and maintaining the Ca^2+^ gradient. This raises the question as to whether LPLs affect acidification and Ca^2+^ transport in a similar way. Proteomic studies have shown that Vcx1 physically interacts with the V_O_ subunits Vma6 and Vph1, as well as the V_1_ subunits Vma1, Vma2, Vma4, Vma10 and Vma13 (Khan et al., 2022; Michaelis et al., 2023). We found that LPC 14:0 blocks Ca^2+^ when added at T=0 min and not at T=10 min. This correlates with blocked acidification when LPC 14:0 is added at T=0 min and not T= 500 s. Similar effects are seen with LPC 18:0 and LPC 18:1, albeit to different degrees. These outcomes correlate with the role of Vcx1 Ca^2+^/H^+^ transport; however, it is not universal for all LPCs tested. When looking at LPC 16:0, we saw that it did not affect Ca^2+^ uptake, yet it blocks acidification. A similar relationship occurs when LPC 16:0 is added later in the reactions. Another inconsistency is seen with LPA, which has no effect on Ca^2+^ or H^+^ transport when added at T=0. When added late, LPA can inhibit Ca^2+^ release and yet it has no effect on vacuole acidification. The lack of an effect on acidification could be because Ca^2+^ did not accumulate outside the vacuole and instead was retained in the lumen. These data suggest that the effects of LPLs on H^+^/Ca^2+^ transport are not linked.

In conclusion, this study shows that vacuole fusion and ion transport can be differentially affected by LPCs depending on acyl chain length and to a lesser extent monounsaturation. We also report that headgroup type influences how LPLs affect fusion if they can also induce positive curvature.

## Data Availability

Correspondence and requests for materials should be addressed to Rutilio Fratti

## Author Contribution

C.Z., R.A.F., designed the research; C.Z., Y.F. J.D.C, A.B., R.A, C.K., V.S., D.G. E.F. J.M.K. R.A.F., obtained the data; C.Z., R.A.F., formal analysis; C.Z., R.A.F., writing original draft; All authors reviewed and edited the manuscript; R.A.F., funding acquisition, supervision, project administration, resources.

## Generative AI statement

The authors declare that generative AI was not used in the writing of this manuscript.

## Supplemental Information

**Supplemental Figure 1.**
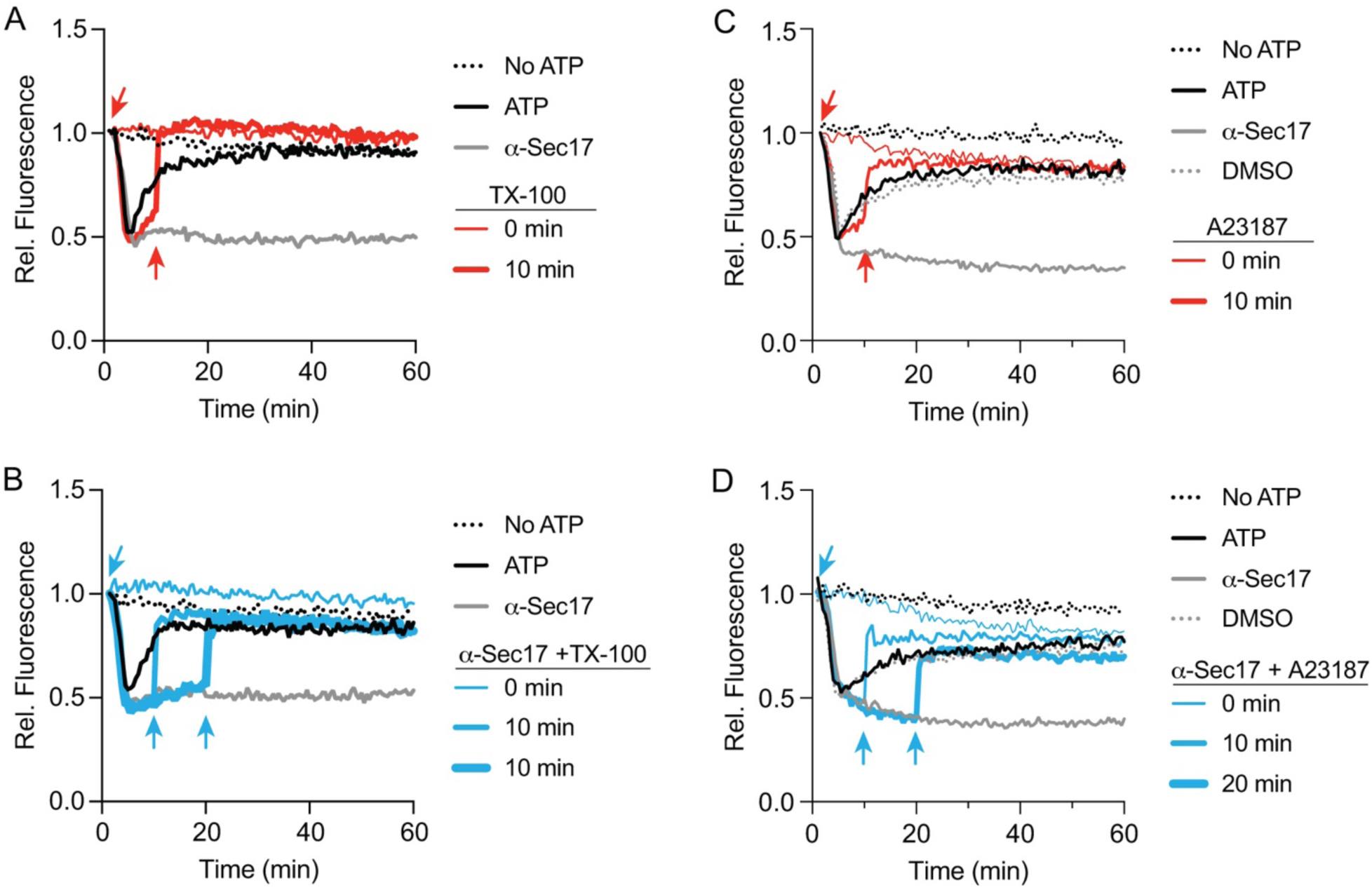
Effect of TX-100 and A23187 on Ca^2+^ transport. Vacuoles were isolated from BJ3505 were treated with 140 µg/mL α-Sec17 IgG, 1% TX-100, 100 µM A23187 or PS added at T=0 or 10 min in the presence or absence of ARS and in the presence of Cal520. **(A)** 1% TX-100 was added at T=0 or 10 min (red arrows). **(B)** 1% TX-100 was added to reactions containing anti-Sec17 IgG at T=0 or 10 min (blue arrows). **(C)** 100 µM A23187 was added at T=0 or 10 min (red arrows). **(B)** 100 µM A23187 was added to reactions containing anti-Sec17 IgG at T=0 or 10 min.

**Supplemental Figure 2.**
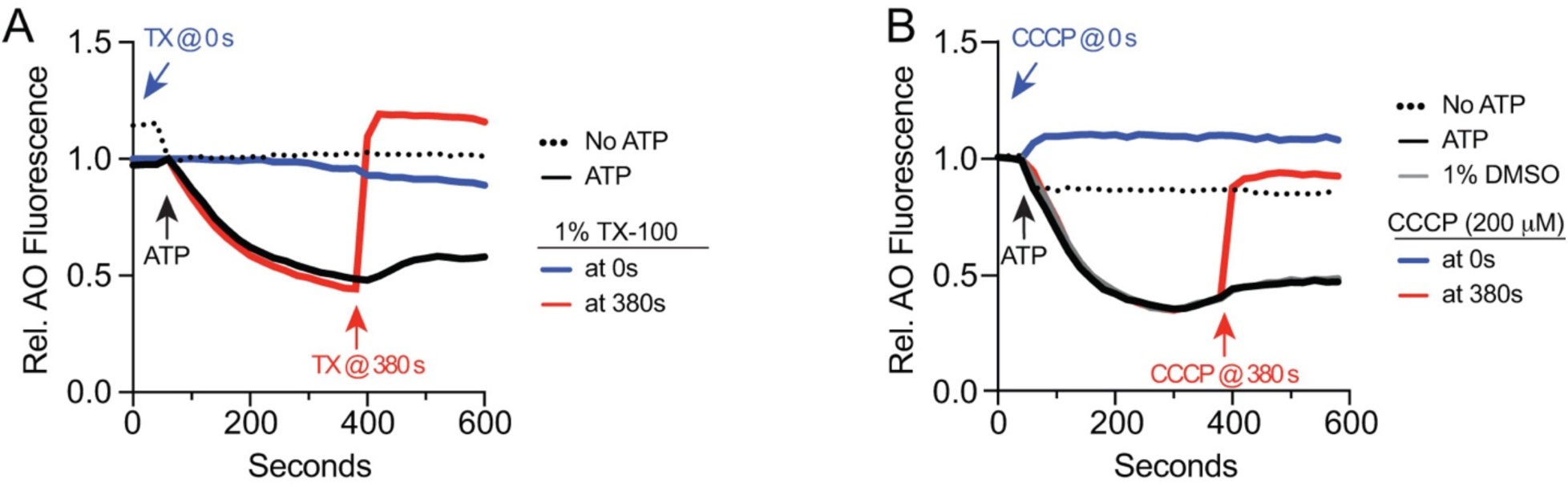
Effect of TX-100 or CCCP on vacuole acidification. BJ3505 vacuoles fusion reactions (2X) were incubated with 1% TX-100 **(A)**, 200 µM CCCP **(B)** or reaction buffer at T=0 or 380 sec in the presence or absence of ARS and in the presence of AO. After 60 s of incubation received ARS or additional buffer and further incubated for a total of 800 s. AO fluorescence was normalized to the fluorescence at the time of adding ARS and set to 1.

